# Long-term imaging reveals behavioral plasticity during *C. elegans* dauer exit

**DOI:** 10.1101/2022.04.25.489360

**Authors:** Friedrich Preusser, Anika Neuschulz, Jan Philipp Junker, Nikolaus Rajewsky, Stephan Preibisch

**Affiliations:** Berlin Institute for Medical Systems Biology (BIMSB), Max Delbrück Center for Molecular Medicine in the Helmholtz Association (MDC), 13125 Berlin, Germany; Institute for Biology, Humboldt University of Berlin, 10099 Berlin, Germany; Janelia Research Campus, Howard Hughes Medical Institute, Ashburn, VA 20147, USA

**Keywords:** Behavioral imaging, *C. elegans* dauer, neuroplasticity

## Abstract

During their lifetime, animals must adapt their behavior to survive in changing environments. This ability requires the nervous system to adjust through dynamic expression of neurotransmitters and receptors but also through growth, spatial reorganization and connectivity while integrating external stimuli. For instance, despite having a fixed neuronal cell lineage, the nematode *Caenorhabditis elegans’* nervous system remains plastic throughout its development. Here, we focus on a specific example of nervous system plasticity, the *C. elegans* dauer exit decision. Under unfavorable conditions, larvae will enter the non-feeding and non-reproductive dauer stage and adapt their behavior to cope with a new environment. Upon improved conditions, this stress resistant developmental stage is actively reversed to resume reproductive development. However, how different environmental stimuli regulate the exit decision mechanism and thereby drive the larva’s behavioral change is unknown. To fill this gap, we developed a new open hardware method for long-term imaging (12h) of *C. elegans* larvae. We identified dauer-specific behavioral motifs and characterized the behavioral trajectory of dauer exit in different environments to identify key decision points. Combining long-term behavioral imaging with transcriptomics, we find that bacterial ingestion triggers a change in neuropeptide gene expression to establish post-dauer behavior. Taken together, we show how a developing nervous system can robustly integrate environmental changes, activate a developmental switch and adapt the organism’s behavior to a new environment.

## Introduction

Animals adapt their behavior in response to changes in the environment, a phenomenon also referred to as behavioral plasticity (1). However, timescales of behavioral adaptation can vary drastically. While some stimuli elicit a fast and short response (e.g. escape) others will influence behavior over minutes (e.g. foraging) or several hours (e.g. day and night cycles). Therefore, characterizing the effect of the environment on an organism’s behavior requires the integration of multiple temporal scales, comprising both short responses and long-term adaptation processes. Studying the adaptation of neuronal circuits and the resulting behavioral dynamics requires experimental tractability i.e. the possibility to identify, trace and compare neuronal circuits across genetic backgrounds and conditions. While this is challenging in complex mammalian circuits, the nematode *Caenorhabditis elegans* (*C. elegans*) has emerged as a suitable model organism for studying the relationship between nervous system plasticity, sensory inputs and behavior by possessing a stereotyped and fully mapped (2), yet flexible (3) nervous system.

The *C. elegans* dauer diapause is a bona fide example of a developmental switch that requires neuronal plasticity (Figure 1a). When encountering an unfavorable environment, e.g. that is lacking a food source, *C. elegans* L1 larvae can actively deviate from the canonical reproductive life cycle and develop into the so-called dauer larva (4). This alternative developmental stage is particularly stress resistant and geared for survival, notably by possessing a thickened, closed cuticle. Dauer larvae cease the uptake of food and can remain in this dormant state for several months (5). Importantly, once environmental conditions improve, dauer larvae will sense and integrate that changed environment and resume reproductive development. Previously, receptors, signaling molecules and transcription factors involved in dauer formation have been studied extensively (for review see (6) and (7)). At the neuronal level, individual remodeling events specific to the dauer larva have been identified (8–12). However, a quantitative and temporally resolved description of the behavioral adaptation process is missing but needed to contextualize molecular and structural changes. Ultimately, linking behavioral adaptation to specific molecular programs (e.g. changes in gene expression), will improve our understanding how a developing nervous system is able to remain plastic and integrate external stimuli.

**Fig. 1.**
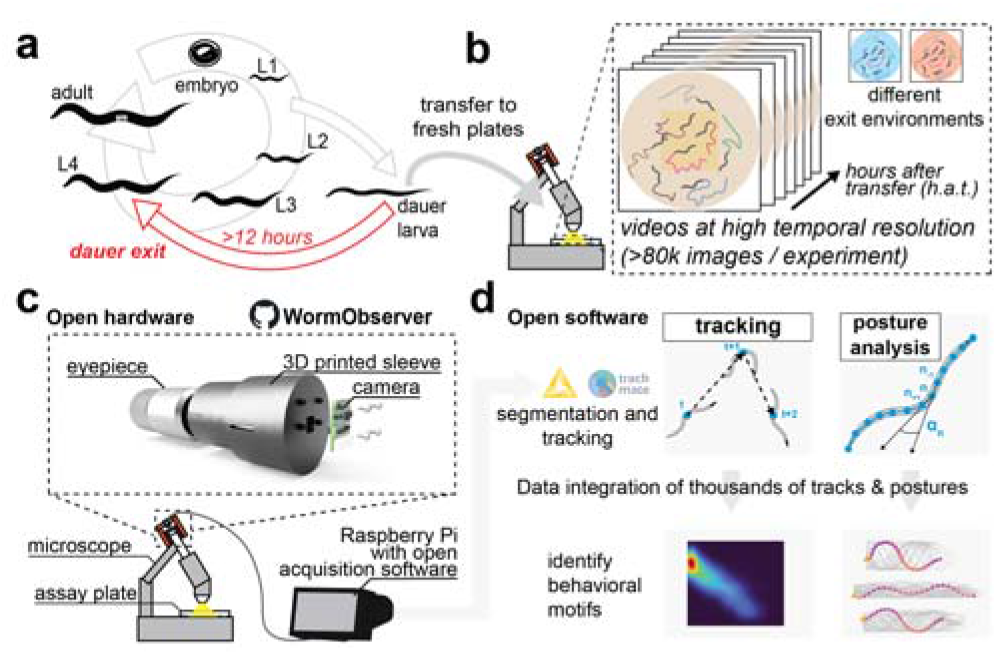
Overview. (a) Dauer exit is a developmental decision to resume reproductive development, which takes more than 12 hours, highlighting the need for long-term behavioral imaging. (b) Overview of the assay used in this study. Dauer larvae are transferred to a new environment (i.e. a fresh plate) and their change in behavior is recorded across several hours. Time resolved measurements will be relative to the time point when worms were transferred to the new environment (hours after transfer or h.a.t.). (c) Sketch of the open hardware; a 3D printed solution for acquiring timelapse videos through the eyepiece of a stereoscope. Camera and acquisition parameters are controlled by a Raspberry Pi, which streams the data to a workstation for data processing. (d) Summary of the software, which integrates KNIME and the TrackMate framework. By integrating data from thousands of tracks and worm postures, users can build robust, quantitative models of *C. elegans* behavior.

Here, we introduce a new open-hardware and opensource platform for quantitative behavioral analysis of *C. elegans.* Although solutions for imaging C. *elegans* behavior have been previously developed (13–17), performing long-term imaging of hundreds of larvae over several hours to extract population dynamics remains challenging. Our WormObserver platform combines long-term population imaging of *C. elegans* larvae and open source analysis software in a modular, end-to-end solution that can be adapted to other research questions (Figure 1b-d). We use the platform to describe the behavioral trajectory of dauer exit and characterize dauer-specific behavioral motifs. Moreover, we identify environmental signals that initiate the exit decision and correlate the resulting changes in behavior with gene expression dynamics.

## Results

### WormObserver: an open hardware solution for long-term behavioral recordings

To capture behavioral changes during the dauer exit decision, a process previously shown to take more than 12 hours (4), we developed the WormObserver, a lightweight, low-cost, open hardware imaging solution that allows parallel tracking of hundreds of worms on culture plates. The WormObserver platform is based on a standard stereoscope, making use of high-quality optical components already available in most laboratories. We designed and 3D printed a two-component adapter that fits the eyepiece of a stereoscope (Figure 1c, Materials and Methods) and holds an attached camera. The camera is connected to a Raspberry Pi, a commercially available, credit-card sized computer, which enables automated video acquisition (18). To provide users with a ready-to-use solution, we developed a graphical user interface (GUI) for easily adapting imaging parameters like acquisition frame rate, resolution and timing of the videos to be acquired. To capture behavioral motifs at high temporal resolution over more than 12 hours, we split the timelapse in short video sequences (8 mins), thereby facilitating tracking and analysis. Acquired videos are streamed directly to a workstation PC applying a custom, open source image processing workflow that uses KNIME (19), ImgLib2 (20) and the TrackMate framework (21) (Materials and Methods). Taken together, combining a 3D printed adapter and off-the shelf hardware components with a standard stereoscope and custom software, we envision our tool to be broadly applicable for many labs, independent of optical or computational expertise.

### Characterizing behavioral adaptation during dauer exit

Dauer larvae have been described to be distinct in their behavioral repertoire and previous studies have improved our understanding of the underlying neuronal plasticity (10, 12, 22, 23). Dauers exhibit increased bouts of prolonged locomotor quiescence but can be highly mobile and are able to nictate (24), a dauer-specific behavior that increases the chances to hitch-hike on other animals to escape an unfavorable environment in natural contexts (25, 26). Moreover, dauers moving on a plate generally appear less bent and stiffer compared to well-fed larvae of the same age (27). Nevertheless, a time-resolved characterization of dauer-exit specific changes in behavior is missing but needed to answer three key questions: First, it is unclear for how long the worm is integrating the changed environment before adapting its behavior. Second, which behavioral motifs are changing during this adaptation process and are all behavioral adaptations occurring at once or in a stepwise manner? And third, are the different environmental stimuli known to regulate dauer exit (i.e. food stimulus, pheromone), all affecting the same or different aspects of behavior? To answer these questions, we focused on the first 12 hours of dauer exit for characterizing behavioral changes and used the condition lacking a food source as a negative control in which dauers cannot successfully exit. We reasoned that behavioral adaptation related to the exit decision would occur before the onset of growth, previously described to resume after ca. 14h of dauer exit (4). We induced the dauer stage by a combination of standardized starvation, crowding and high-temperature conditions (see Materials and Methods). Next, extracted dauer larvae were shifted from starved to fresh plates with or without a bacterial food source and monitored for 12 hours. To compare behavior across conditions, we refer to each timepoint as hours after transfer *(h.a.t.)* to a fresh plate (Figure 1b).

### In a new environment, dauers will switch from behavioral quiescence to a highly mobile state

We first asked the question whether velocity, i.e. the speed by which the worms move on the plate was changing during the course of dauer exit. As expected, dauers that were transferred to a new environment initially exhibited very slow movement or no movement at all, characterized by high angular velocity (i.e. path curvature) and low velocity. The previously described dauer specific locomotion quiescence (22) was observed for dauers in all tested conditions and is independent of the presence of bacteria on the plate (Figure 2a, Supplemental Figure S1). However, many larvae remained highly mobile (high velocity) and moved straight in one direction (low angular velocity). This initial division into a quiescent and a highly mobile regime appeared in all our experiments and is also recapitulated when quantifying behavior of *daf-2* dauer larvae that cannot exit the dauer stage (Figure 2b, Supplemental Figure S1). We therefore conclude that dauer larvae, when put onto fresh plates, remain quiescent but can explore the environment in a specific dispersal mode characterized by fast movement into one direction. This is also consistent with the observation that dauers can be highly mobile (27). Moreover, when extending our analysis beyond the first hours after transfer, we noticed that the quiescent fraction disappeared 3 h.a.t. At this time point, all animals displayed a strong dispersal phenotype characterized by high velocity and low angular velocity (Figure 2c&d). This was similar across the tested conditions, although the exact timing varied, with the notable exception of *daf-2* dauer larvae, where a significant fraction remained in the quiescent phenotype (Figure 2b, Supplemental Figure S1). Taken together, we observe that independent of a bacterial food source, 3h after being exposed to a new environment, wild type dauer larvae are exhibiting a first behavioral switch from quiescence to a highly mobile dispersal-like behavior. Importantly, this first behavioral switch is independent of the presence of a food source and was observed in all tested conditions.

**Fig. 2.**
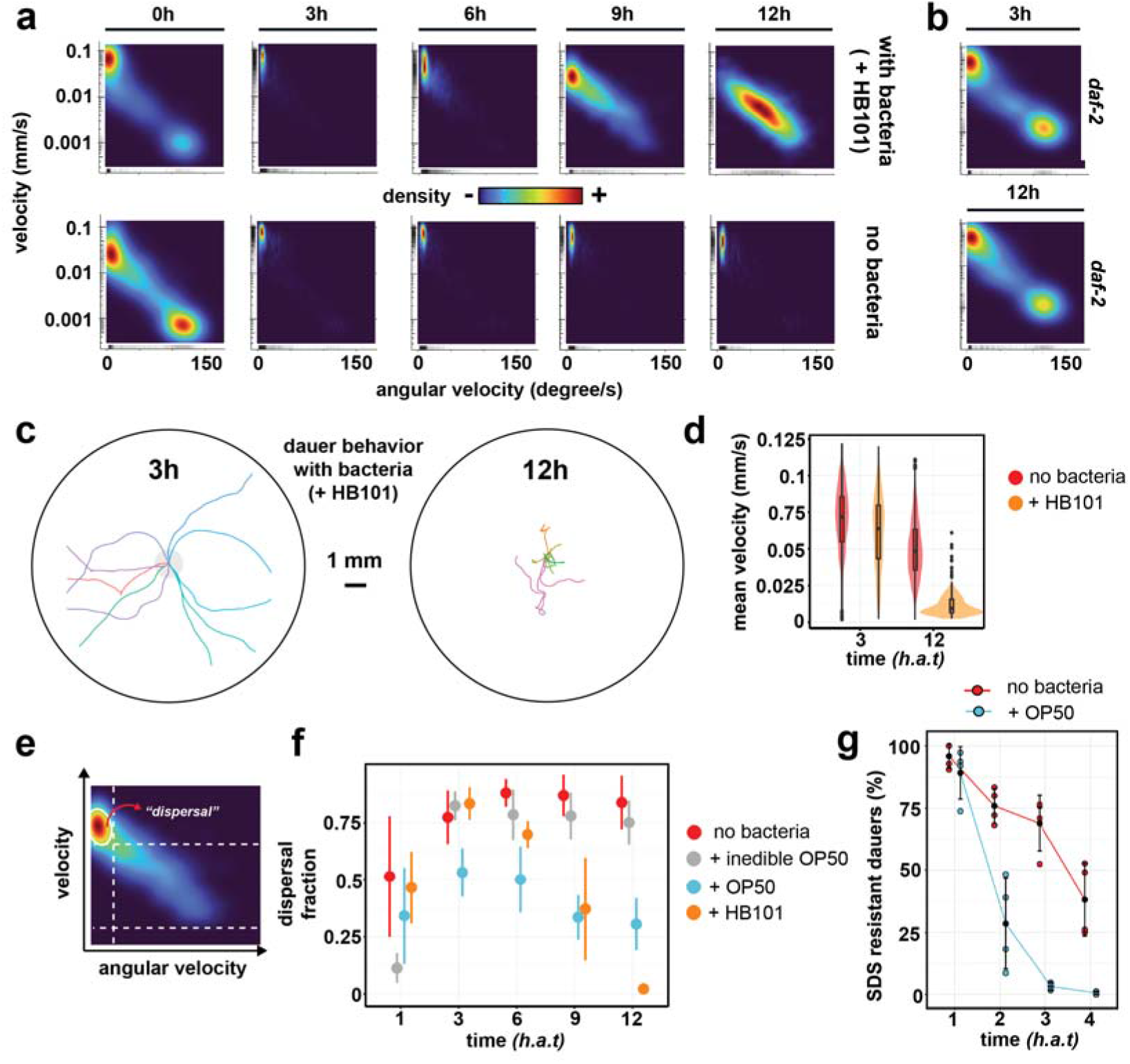
A food-dependent behavioral switch during dauer exit. (a) Two-dimensional probability distributions of speed (mm/s) and angular velocity (degrees/s) averaged over 10 s time windows across the first 12 hours of dauer exit, highlighting the effect of a food source on behavior. Data is from n=3 experiments and each time point contains data from at least 250 tracks (average number of tracks = 1262). (b) Two-dimensional probability distributions obtained from daf-2 dauers, n=2 experiments. (c) 10 example tracks, comparing dauer behavior before and after the food-dependent behavioral switch. (d) Mean velocity (calculated per track) before and after the food-dependent behavioral switch. (e) Schematic summarizing how the dispersal phenotype was defined, based on thresholds for velocity and angular velocity (see Materials and Methods). (f) Relative abundance of dispersal behavior as defined in (e). Error bars depict standard deviation of at least 2 experiments. (g) Percentage of dauers that survive SDS treatment at different time points during dauer exit. This assay can be used as a proxy for inferring onset of pharyngeal pumping. Dots represent experimental replicates.

### A second behavioral switch depends on a food stimulus

When extending our analysis beyond the first three hours, we noticed a second behavioral adaptation that was dependent on the presence of an edible food source. After being exposed to the new environment for approximately 6 hours, larvae that were placed on a high quality food source decreased speed but increased turning frequency, resembling canonical behavior of adult worms (28) (Figure 2a). After 12h, dauer specific dispersal behavior was completely abolished (Figure 2c&d). In contrast, dauers placed on plates lacking food continued moving at high velocity and low angular velocity and did not exit dispersal behavior (Figure 2a&d). When quantifying the relative fraction of dispersal behavior across conditions and timepoints (Figure 2e), we noticed that the behavioral switch from dispersal to dwelling started to occur with high quality (HB101) and low quality food (OP50), in both cases after 6 h.a.t. (Figure 2f). Interestingly, larvae placed on bacteria that were previously treated with an antibiotic, making them too large to be ingested by the worm, did not exit the dauer-specific dispersal behavior (Figure 2f). This finding suggested that the ingestion of bacteria is crucial for inducing the behavioral switch. To further investigate this, we confirmed that onset of pharyngeal pumping happens prior to 6 *h.a.t.,* however with delayed timing if a food source is lacking (Figure 2g) (4). Taken together, we have characterized a second behavioral switch that consists in exiting the dauer-specific dispersal phenotype and that is, in contrast to the first switch, dependent on the presence of a bacterial food source and therefore indicative of successful dauer exit. Furthermore, we note that it is the ingestion (and not the presence) of bacteria that drives this behavioral change, matching the observation that by the time the switch occurs, dauers will have resumed pharyngeal pumping.

### Posture adaptation is independent of the presence of a food source

*C. elegans* posture, i.e. the worm’s shape, has been previously established as a robust measure for inferring behavioral motifs based on environmental stimuli or genetic differences (29–34). Specifically, we hypothesized that the dauer-specific stiff and more narrow posture would disappear during dauer exit. Moreover, we asked the question whether the dauer-specific dispersal phenotype would be reflected at the posture level. As previously described for adult worms (29), segmented worm shapes extracted from moving worms across all timepoints have low complexity and can be explained by a combination of eigenvectors obtained from the covariance matrix of intersegment angles, so-called “Eigenworms” (Supplemental Figure S2). We created a dauer-specific posture library (i.e. a postural syntax) for each condition (34) and plotted relative posture abundances across all timepoints (Figure 3a, Materials and Methods). Posture abundances during dauer exit were consistent across experimental repeats (Supplemental Figure S3). Interestingly, we detected a shift in postural syntax at about 6 *h.a.t* when dauers were shifted to a bacterial food source (Figure 3b). We next aimed at further characterizing postural syntax dynamics across conditions and performed a second round of clustering on the previously defined posture libraries to identify the 3 most diverging posture groups (i.e. meta postures, p^1^, p^2^, p^3^) and compare their relative abundance across conditions (Supplemental Figure S4). While skeletons of the first meta posture p^1^ resemble the dauer-specific constricted posture, p^2^ and p^3^ contain more bended postures (Figure 3c). Consistently, intersegment angle distributions differ significantly between meta postures (Figure 3d). Next, we quantified the relative abundance of the dauer-specific p^1^ postures across timepoints and noticed a decrease in relative abundance within the first 6 *h.a.t.* in all tested conditions except for *daf-2* mutants that exhibited an impaired transition (Figure 3e and Supplemental Figure S5). It is important to note that due to our skeletonization implementation (Materials and Methods), our posture libraries did not contain overlapping postures of turning worms. Hence, the observed decrease in dauer specific postures could be the result of an increase of omega turn frequency as observed in starved worms (35). To test this possibility, we trained a random forest classification model to robustly identify omega turns (Supplemental Figure S6, Materials and Methods) and quantified the fraction of turning worms during dauer exit. While the fraction of omega turns increases in the +OP50 condition, omega turn frequency remains low when a bacterial food source is lacking (Figure 3f). Thus, we conclude that the measured posture switch detected in conditions with and without a bacterial food source, cannot be solely explained by an increase in turn frequency. Taken together, we detect a new example of behavioral adaptation that consists in switching from a dauer-specific stiff and thin posture to a more bended, canonical *C. elegans* posture. This change in postural syntax happens within the first 6 *h.a.t.*, but is independent of the presence of a food source, does not explain the dauer-specific dispersal behavior and might be driven by other environmental factors (e.g. pheromone concentration).

**Fig. 3.**
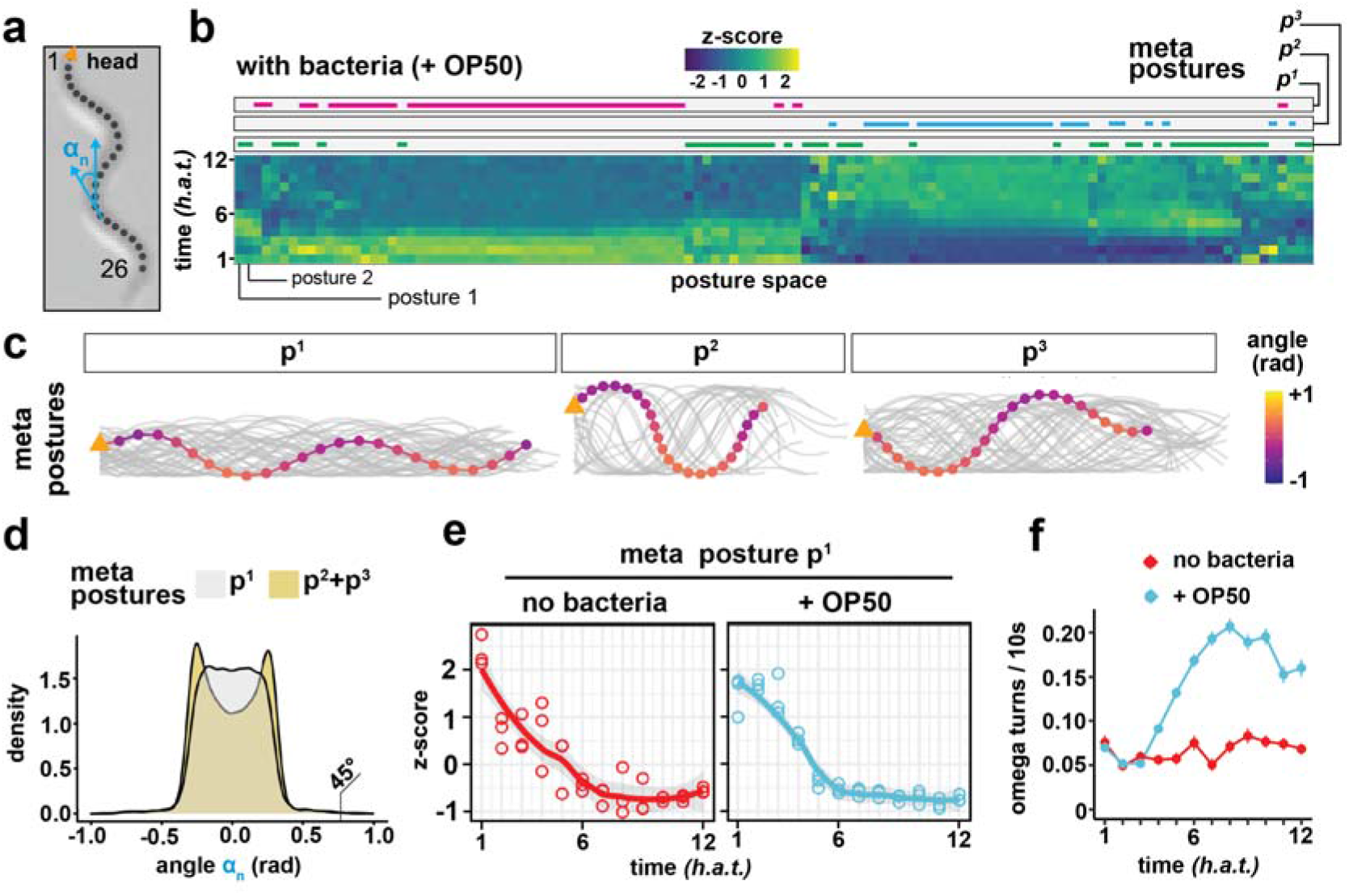
Postural adaptation during dauer exit. (a) Schematic of a worm skeleton. For each skeleton, we calculate a vector of 24 intersegment angles that describe the worm’s posture. (b) Heatmap of the relative occurrences of 120 postures during dauer exit on a food source as quantified from n=3 experiments. Color indicates whether a posture is over or underrepresented at that time point relative to all time points. The upper bars indicate three meta posture groups, each comprising several postures, and their allocation within the heatmap. Meta postures were obtained by a second k-means clustering with k=3 of all postures from all experimental conditions. (c) The three meta postures that were calculated across all conditions. All postures of one group are depicted in gray and one example posture is shown in color. (d) Distribution of all intersegment angles, calculated as shown in (a). The dauer-specific meta posture p1 is compared to the two other meta postures p2 and p3. p1 contains 46 postures, p2 and p3 contain 28 and 46 postures, respectively. (e) Relative occurrence of the meta posture p1 during dauer exit, normalized to all time points of the depicted experimental condition. Each dot represents one experiment of n=3 experiments. (f) Omega turn frequency during dauer exit, quantified with a custom random forest classifier (see Materials and Methods). Depicted is the mean, error bars are SEM.

### Genes driving behavioral changes during dauer exit

Next, we asked the question whether the two main behavioral switches quantified by imaging (Figure 4a) would also be reflected at the level of gene expression, potentially through the expression of specific neuropeptides, receptors and ion channels in neurons, previously described to be sufficient to cause drastic behavioral changes in *C. elegans* (36–38) and dauers specifically (10, 39). For instance, insulin-like peptides (ILPs), which are expressed in the head of the worm (40), have been previously shown to be implicated in regulating dauer arrest (40–42). We therefore performed a low-input RNA sequencing method (43) on *C. elegans* heads specifically at 4 different timepoints during dauer exit, exposing recovering dauers to an environment with or without a food source, thereby mirroring the conditions used for behavioral analysis (Figure 4b, Materials and Methods). Next, we asked the question whether the behavioral switches were recapitulated at the level of the transcriptome by plotting gene expression in PCA space across all conditions and timepoints. Strikingly, for the condition lacking a food source, head-specific transcriptomes of 3h, 6h and 9h timepoints were similar as indicated by PCA clustering (Figure 4c). Analogous to the behavioral data, if bacteria were provided, the timepoints 6h and 9h separated as well (Figure 4d), indicating a two-step process driving the exit decision. We next wanted to characterize both behavioral adaptation switches in greater detail by identifying differentially expressed genes between the two conditions at different timepoints. Here, we focus on transcripts that are relevant for the behavioral adaptation process, but we offer an interactive online resource (https://www.bit.ly/dauer_exit) for exploring the whole dataset. To characterize differential expression during the first food-independent behavioral switch, we computed a list of genes that were differentially expressed in both conditions when comparing the 0h and 3h timepoints. This revealed a list of genes, which contained all members of the arcasoid beta-oxidation pathway (44), all being downregulated in the first 3 *h.a.t.* (Supplemental Figure S7). Additionally, we noticed significant upregulation of the insulin-like peptide DAF-28, described to be transcriptionally repressed by increased pheromone concentration (42, 45) and required for dauer exit (46). Taken together, we note that early in dauer exit, potentially driven by changes in pheromone concentration (e.g. through downregulated beta-oxidation pathway activity), DAF-28 expression goes up. This change in DAF-28 expression is independent of the presence of a food source and therefore indicative of an early, intermediate behavioral state that takes place while the exit process is still ongoing. Next, we sought to investigate the second food-dependent behavioral switch that depends on a bacterial food source, which was already apparent at the transcriptome (Figure 4c). To identify biological pathways that are differentially regulated between with and without bacteria conditions, we performed functional enrichment analysis using differentially expressed genes as input (Figure 4e, Materials and Methods). At 6 h.a.t, coinciding with the second behavioral switch to post-dauer behavior, the most significantly differentially expressed gene ontology (GO) - term was “neuropeptide signaling pathway” (p-value: 1.66×10-8) (Figure 3f). The differentially expressed genes within this GO-term revealed numerous neuropeptides, comprising both FMRFamide peptides (FLPs) and neuropeptide-like proteins (NLPs), previously described to modulate behavior in *C. elegans* (47) as well as some of the corresponding receptors (Supplemental Figure S7). We found that almost all (23/24) neuropeptides were downregulated during dauer exit. Strikingly, one neuropeptide, NLP-24, was upregulated (Figure 4f). Besides NLPs and FLPs, we also note differential regulation of ILPs, the third class of neuropeptides in C. elegans, at 6 h.a.t when comparing environmental conditions (Supplemental Figure S7). Consistent with insulin signaling driving successful dauer exit, INS-17, an insulin antagonist (48) is downregulated while the counteracting INS-4, a DAF-2 agonist (45), is upregulated. In summary, we find that dauers that have been placed in an environment containing a food source will exhibit a behavioral change after 6 *h.a.t.,* which coincides with a drastic change in neuropeptide gene expression.

**Fig. 4.**
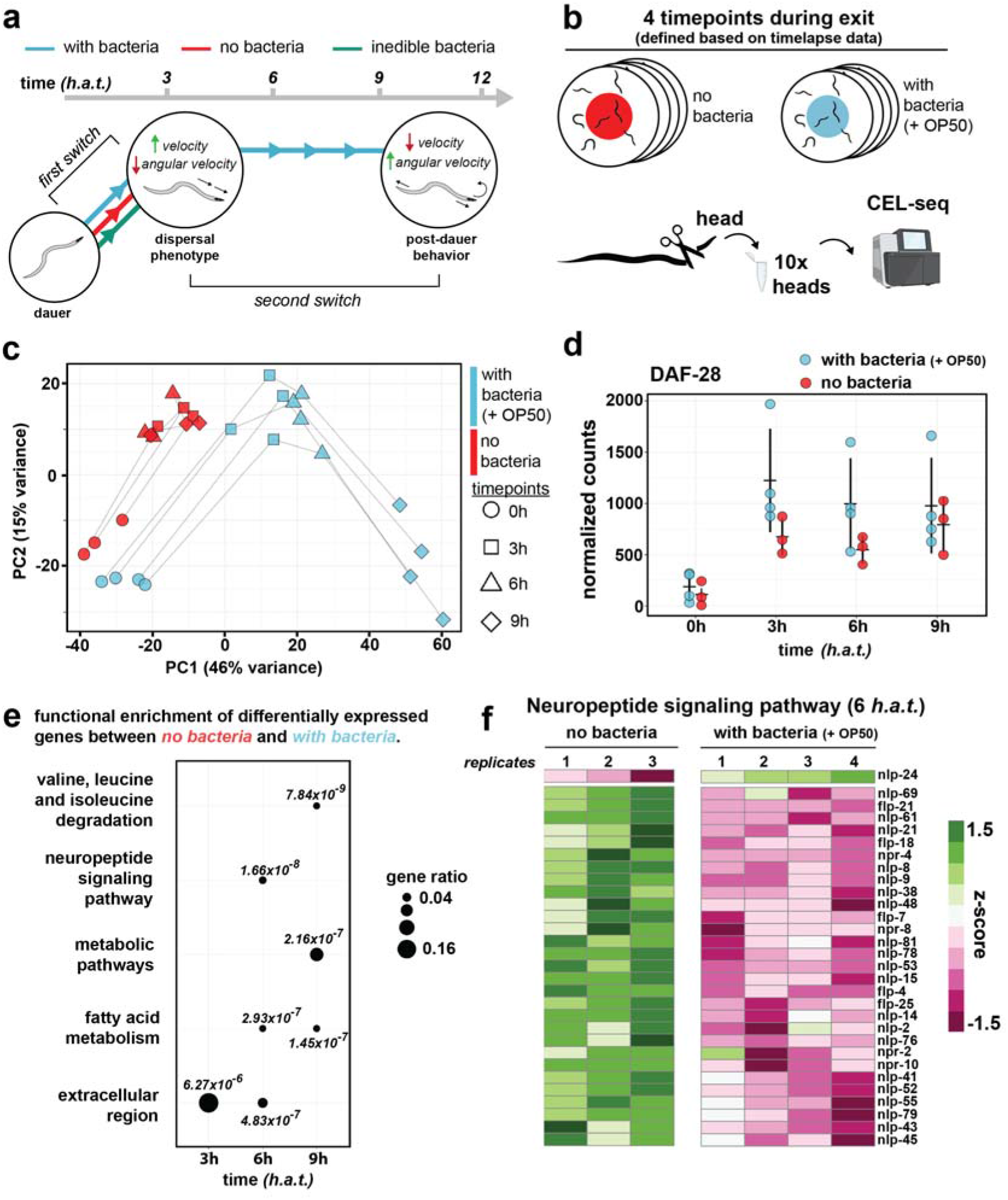
Transcriptome profiling reveals dynamic neuropeptide signaling during dauer exit. (a) Schematic summarizing the two main behavioral switches identified by behavioral imaging. (b) Schematic of the head-specific low-input mRNA sequencing method. For each timepoint and each condition, 10 heads were sequenced. (c) PCA of all samples, using the log transformed and normalized counts of the 500 most variable genes as input. (d) Normalized counts of DAF-28 across both conditions and all time points. Depicted is the mean +/- standard deviation. (e) Dotplot depicting the result of the GO-term analysis. For each time point, differentially expressed genes between the two conditions were determined and the result was used for GO-term analysis. Size of the dot scales with gene ratio. Gene ratio refers to the proportion of genes in the input list that are annotated to that GO-term. Shown are the top 3 GO-terms for each time point and p-values are indicated next to each dot. (f) Heatmap of genes contained within the “neuropeptide signaling pathway” GO-term, comparing both conditions at 6 *h.a.t.* One column shows one experimental repeat.

## Discussion

In this work, we present the WormObserver as a new, easy-to-use and cost-effective open hardware solution for the analysis of *C. elegans* behavior. We envision our method to be useful for many labs independent of computational expertise to perform long-term acquisitions and analysis of *C. elegans* behavior. A description of the workflow, including detailed instructions can be accessed under https://github.com/Fritze/WormObserver. The modular nature of our system allows for the integration of more advanced opto-electronics or software components (e.g. optogenetics (49) or other tracking software (50, 51)). Moreover, given its cost (<150$) and straightforward assembly, the platform can be considered for larger screens involving several microscopes running in parallel. We also note that in the future, one might consider running parts of the analysis directly at the microscope to reduce data footprint and analysis time, for instance by only saving worm centroid locations. As cost-effective GPU-equipped minicomputers similar to the RaspberryPi are emerging (e.g. NVIDIA Jetson), application of custom deep learning models to perform these tasks directly at the microscope becomes feasible. Using the WormObserver, our behavioral analysis of dauer exit revealed two behavioral adaptations that have not been described previously:

1. When placed into a new environment, dauer larvae will exit locomotion quiescence and move into a highly mobile, dispersal state. In parallel, dauers will abandon the dauer-specific narrow posture. This first behavioral adaptation happens within 6 h.a.t and is independent of the presence of food. In the wild, dauers can overcome long distances and disperse to more favorable environments by phoresy, the ability to be carried on by another animal (25, 26). However, an intermediate, highly mobile behavioral state occurring early in the exit decision process, would enable fast dispersal to more favorable environments across smaller distances. This hypothesis is supported by the fact that *C. elegans’* “boom-and-bust” lifecycle in the wild crucially depends on active migration during the dauer stage (52). Interestingly, dauers exhibit distinct alterations in muscle structure, notably increased muscle arm extension, which may facilitate dispersion (53).In summary, we hypothesize that the ability to switch to a highly mobile dispersal phenotype is likely to have evolved as an additional behavioral strategy to increase the dauer larva’s ability to efficiently disperse to more favorable environments. Strikingly, co-occurring with this first behavioral adaptation, we notice the upregulation of the only insulin-like peptide that is required for dauer exit, DAF-28. With DAF-28 being expressed in ASJ neurons (42), the critical sensory neuron for dauer exit (54), it will be interesting to further explore its role in promoting behavioral states during dauer exit.
2. If the new environment contains a food source, at 6 *h. a.t.*, dauers, which have resumed pharyngeal pumping at this point, start to ingest bacteria and eventually leave the transient dispersal-like state to adapt post-dauer behavior. Strikingly, we find that the two transitions observed at the behavioral level are both reflected at the transcriptome level as well. We also discovered that regulation of neuropeptides in response to food drives the dauer exit decision and initiates the final behavioral switch. Neuropeptides are well conserved, short amino acid sequences that can directly act as neurotransmitters but also have neuromodulator capacity (55). In *C. elegans,* the dynamic expression of neuropeptides has been described to control several fundamental behavioral programs, e.g. mating (56), feeding (38), sleep (57) and others (47). More specifically, in the context of dauer larvae, previous work has identified increased neuropeptide signaling as an important neuromodulatory event for promoting the dauer entry decision (58, 59). Hence, adjusting the neural transcriptome at the level of diffusible signaling-molecules, i. e. neuropeptides, increases the larva’s behavioral repertoire. However, we also find that some neuropeptides are specifically upregulated during dauer exit on a food source (e.g. NLP-24, INS-4), highlighting the complexity of neuropeptide signaling. Given its previously reported role in mediating pharyngeal pumping in starved worms (60), upregulation of NLP-24 could regulate onset of pharyngeal pumping during dauer exit but further experiments will be needed to confirm this hypothesis. Previously, NAD+, which is contained within the bacterial food source, has been shown to promote pharyngeal pumping in dauer-exiting larvae (61). Here, we show that only ingestible bacteria will later in the process promote another behavioral switch affecting locomotion. It will be interesting to further characterize which components of the ingested bacteria are required for this process and how these components are sensed and processed. Finally, we note that in the future, resolving transcriptional dynamics of these neuropeptides at the level of individual neurons (e.g. through single-cell RNA sequencing (62, 63)) will further improve our understanding of the underlying circuit adaptations. In summary, while comprehensive studies of the combined effect of neuropeptide gene expression changes are missing, our study sheds light on the role of neuropeptides in establishing a developmental decision in response to environmental stimuli. By correlating differential expression with longterm behavior data obtained in the same conditions, we associate gene expression changes to behavioral switches during dauer exit, making a first step towards answering the question how nervous system plasticity can be achieved by dynamic expression of neuromodulators. Thus, our work also highlights how the combination of long-term behavioral imaging and gene expression analysis can improve our understanding of developmental and neuronal plasticity.

## Code availability

A full description of the WormObserver, including detailed instructions for all hardware and software components can be found at: https://github.com/Fritze/WormObserver All code used for analysis of tracking and sequencing data is available at: https://github.com/Fritze/WormObserver/tree/master/code

## Data availability

Raw data for all imaging experiments can be accessed under: https://bit.ly/dauer_exit_image_data

Sequencing data is available under: https://bit.ly/dauer_exit_sequencing_data

RNA sequencing results can be interactively explored at: https://www.bit.ly/dauer_exit

## ACKNOWLEDGEMENTS

We acknowledge Andrew Woehler and the Systems Biology Imaging Facility for providing access to their microscopes. We thank Mandy Terne for excellent technical assistance, Asija Diag for guidance regarding worm cutting, and Jonathan Fröhlich for critical feedback on the manuscript. We also thank the Caenorhabditis Genetics Center at the University of Minnesota for providing C. elegans strains. F.P. was funded by a PhD fellowship from Studienstiftung des deutschen Volkes and MDC-Berlin. A.N. was funded by a PhD fellowship from Studienstiftung des deutschen Volkes. Work in J.P.J.’s group was funded by ERC starting grant SPACEVAR. S.P. was funded by MDC Berlin and HHMI Janelia. F.P. and S.P. were supported by HFSP grant RGP0021/2018-102.

## AUTHOR CONTRIBUTIONS

F.P., N.R. and S.P. conceived the project. F.P. performed all experiments, analyzed the data and wrote the manuscript. A.N. and J.P.J. contributed expertise for RNA sequencing experiments and A.N. generated count matrices. S.P. supervised the work and contributed with resources, project administration and funding acquisition.

## Materials and Methods

### C.elegans strains and handling

Worms were maintained at 15°C on nematode growth medium (NGM) with Escherichia Coli OP50 as food source as previously described. Dauer formation was induced by transferring a mixed population of ca. 50-100 worms to 6cm NGM plates seeded with E. coli OP50. Plates were then incubated for at least 7 days at 25°C. After 7-10 days, worms were washed off with M9, washed twice to eliminate remaining bacteria and 1% SDS solution was added. The mixture was incubated for 30 mins at room temperature (4). After an additional washing step with M9 to eliminate SDS, worms were transferred to an unseeded NGM plate. After 30 mins, once the drop of M9 dried out, dauers, which can survive SDS treatment (unlike all other *C. elegans* stages), crawled out of the mix of dead worms and were collected. All experiments were carried out with the N2 Bristol strain (obtained from CGC) except for experiments involving a daf-2 insulin mutant. Animals carrying the *daf-2* (e1370) mutation were incubated at 25°C for at least 7 days to obtain pure dauers and subjected to the same SDS-selection procedure prior to imaging.

### Assay plates

We used commercially available freeze-dried E.coli OP50 (+OP50, available from LabTIE) or HB101 (+HB101, available from CGC) bacteria for conditions involving a bacterial food source. Dead OP50 bacteria were prepared according to the manufacturer’s instructions. HB101 were cultured overnight at 37°C. Both cultures were diluted 1:10, added to 3cm NGM plates, UV irradiated to abolish bacterial growth and kept for storage at 4°C. Inedible bacteria were prepared by incubating E.coli OP50-GFP (available from CGC) with Aztreonam (Cayman Chemical, CAY19784-1). The resulting elongated bacteria were visually inspected under a fluorescent microscope, seeded on NGM medium, UV irradiated and kept for storage at 4°C. For the “no bacteria” condition plates contained only 2% Agar.

### Pharyngeal pumping assay

N2 dauers were obtained as described above and placed on assay plates with or without bacteria (see above). Every hour, ca. 50-100 worms were washed once in M9, added to 1% SDS and incubated for 15 minutes at room temperature. Next, an additional wash in M9 was performed and worms were checked for viability on an unseeded NGM plate.

### WormObserver and imaging

For each imaging experiment ca. 250 dauers were transferred in a small drop of M9 and added to 3cm plates (for preparation see above). We used a Zeiss Discovery V8 stereoscope with illumination from below the plate and a Plan S 1x objective with 3.2x zoom. To filter out blue light from the illumination source, a red filter (Edmon Optics, 53-699) was placed in between the light source and the assay plate. The WormObserver hardware consists of two parts: A custom 3D-printed eyepiece sleeve with a fixed 8 Megapixel Raspberry Pi camera and an attached Raspberry Pi model 3 with a touchscreen display for defining acquisition parameters. We provide all details for fabricating the hardware components together with the required software and descriptions under https://github.com/Fritze/WormObserver/. For all experiments shown here, we recorded at least 100 consecutive videos of 8 minutes (>13 hours) and a framerate of 5 fps. All timelapse data was acquired at room temperature (20°C). Experimental repeats were performed on different days.

### Imaging and tracking

The custom, open-source KNIME workflow that is designed for automated processing of the data can be assessed under https://github.com/Fritze/WormObserver/tree/master/KNIME.

Users only need to define the location of the incoming videos as sent by the RaspberryPi and all image processing steps are performed automatically with no user intervention needed. H264-compressed videos are automatically converted into images using ffmpeg and downsampled to 2 fps for tracking. Briefly, worms are segmented after background subtraction, filtering and size selection of segmented connected components. Tracks are identified using the TrackMate (21) node and worm skeletons extracted using the “skeletonize” ImageJ command in KNIME. In summary, each video of a given timelapse experiment is summarized in a result table containing an ID, the relative position, track statistics, omega turn prediction score, the segmented shape and the skeleton for all detected and tracked worms. A workstation (Intel 2.6 GHz CPU, 64 GB RAM) was used for processing all datasets. Processing 100 timepoints took about 17h per dataset, which is only 4h longer than the runtime of the timelapse, highlighting the value of parallelizing acquisition and analysis for long timelapse experiments.

### Tracking data analysis

We wrote custom code in R to process result files as given out by the KNIME workflow. All analysis steps presented in this paper can be performed by calling R-scripts from the command line. All scripts can be accessed under https://github.com/Fritze/WormObserver/tree/master/code/tracking.

For all analysis, we discard the first 30 mins of the timelapse to avoid artifacts introduced by worm handling. Briefly, we calculate velocity and angular velocity per track over 10s bins. Only tracks without gaps and of minimum length of 20 frames (i.e. 10 seconds at 2 fps) were considered. We additionally exclude non-moving tracks to avoid segmentation artifacts. Dispersal behavior was defined as tracks having a minimum velocity of 0.01 mm/s and angular velocity below 15 °/s.

### Skeletonization

For every frame and segmented worm, a skeleton was extracted and saved from within the KNIME workflow. All subsequent analysis steps were performed in R and relevant source code can be accessed under https://github.com/Fritze/WormObserver/tree/master/code/tracking/. Briefly, for each skeleton we first estimate the worm’s head position based on the direction of movement. Next, only skeletons that can be unambiguously identified (i.e. no branches) and of a minimum length of 26 pixels are retained. Finally, for skeletons where the first step could not assign head position on the direction of movement, information from adjacent frames was used. If adjacent frames didn’t contain head position information, the skeleton was discarded. We manually confirmed this approach to be a good conservative estimate of the worm’s skeleton and head position. For calculating Eigenworms, we focused on moving worms and only included tracks with a minimum track duration of 5 seconds, velocity > 0.01 mm/s and angular velocity < 50°/s.

### Posture analysis

For each skeleton, we then fit 26 equally spaced segments on the midline of the worm and calculate a vector of 24 intersegment angles, previously shown to be sufficient for describing C. elegans posture (32)). To cluster postures based on intersegment angles obtained after skeletonization, we used an approach adapted from (34). For this analysis, we only considered tracks that were successfully tracked and skeletonized for at least 5 seconds without interruption. Since we do not follow individual worms and therefore image at lower magnification to capture a larger number of larvae within the field of view, we could not distinguish dorsal and ventral turns in our data. After clustering with k=200, we next identified mirrored posture pairs and treated each pair as if it was the same posture for further analysis. This reduced the complexity of our posture library by ca. 50% depending on the analyzed condition. This first round of clustering was done individually for each condition but across experimental repeats. To compare relative abundances of certain posture types (i.e. meta postures) across conditions, we subjected all previously identified postures to a second round of k-means clustering (k=3) and plotted the relative abundance of the three meta postures across conditions and experimental repeats. We note that the resulting posture libraries did not contain any self-occluded shapes (i.e. omega turns) as these did not pass our criteria for successful skeletonization. To be able to measure the abundance of omega turns within our datasets across timepoints and conditions, we used 17 image features extracted from segmented worm bitmasks and trained a random forest model for detecting omega turns. The classification is integrated within the image processing workflow in KNIME and flags every frame of a track with either 0 or 1 if an omega shape was detected. Next, we bin all omega turn occurrences over a period of 10s to avoid over counting still animals (i.e. if more than one frame is flagged as turn within the 10s window, it will be counted only once).

### RNA-seq - library preparation

N2 dauer larvae were cultured and extracted as described above. Assay conditions were the same as for the timelapse imaging experiments and both conditions (no bacteria and + OP50) were processed in parallel. Every hour, a fraction of dauers was washed off the assay plates and anesthetized with 1mM tetramisole. Under a dissecting microscope, the worm’s head was cut off just below the terminal bulb of the pharynx. For each timepoint 10 heads were pooled into a pre-cooled Eppendorf tube on dry ice, containing 500 μl trizol and 1 μl GlycoBlue. Samples were frozen at −80°C for storage and later subjected to RNA sequencing according to the Tomo-Seq protocol (64) with minor modifications. Briefly, after thawing, samples were mixed thoroughly and incubated for 5 min at room temperature. Subsequently, 100 μl of chloroform were added to each sample, mixed well and incubated for 5 min. Samples were then centrifuged at 12,000 g for 15 min at 4°C. Next, the aqueous phase was carefully transferred to an Eppendorf LoBind containing 250 μl isopropanol and precipitated overnight at −20°C. Samples were centrifuged for 10 min at 12,000 g and at 4°C. Next, the supernatant was removed and the RNA pellet was washed with 75% freshly prepared ethanol. After washing, the pellet was dried and, once completely dry, resuspended in 2 μl molecular grade water. 1 μl of the sample was transferred to a fresh low binding PCR tube containing barcoded oligo(dT) primer. Reverse transcription was performed with the Superscript II kit according to (Holler and Junker, 2019). First strand synthesis was performed for 1 h at 42°C. Second strand synthesis was performed for 1 h at 16°C. Next, samples were pooled and purified with Agencourt AMPureXP beads. IVT was performed for 16h at 37°C. After fragmentation and purification, RNA was reverse transcribed with superscript II and amplified by PCR. Concentration and size distribution of the resulting cDNA were measured with Qubit and Tapestation. mRNA Libraries were sequenced on a NextSeq500 in paired end mode with 150 cycles.

### RNA-seq - Differential expression analysis

Raw sequencing basecalls were demultiplexed and converted to FASTQ format using custom python scripts. RNA-seq reads were mapped to the ce11/WBcel235 genome assembly using STAR_2.7.3a (65), annotated using Rsubread (66) and count matrices were generated using custom python scripts. PCA was done on normalized log transformed read counts of the 500 most variable genes across conditions and timepoints. Differential expression changes were determined with DESeq2 (67), with lfc shrinkage turned on. Differentially expressed genes were called with a significance cut-off of 0.05 (p-value adjusted) and a minimum log2 fold change of 1 or −1. For GO term analysis, the gprofiler2 package (68) was used with a significance cut-off of p-value 0.01 and a custom background model that contained all genes that we detected with our method across all conditions and timepoints with at least 10 reads. The dataset can be interactively explored under https://www.bit.ly/dauer_exit.

## Supplemental figures

**Supplemental figure S1.**
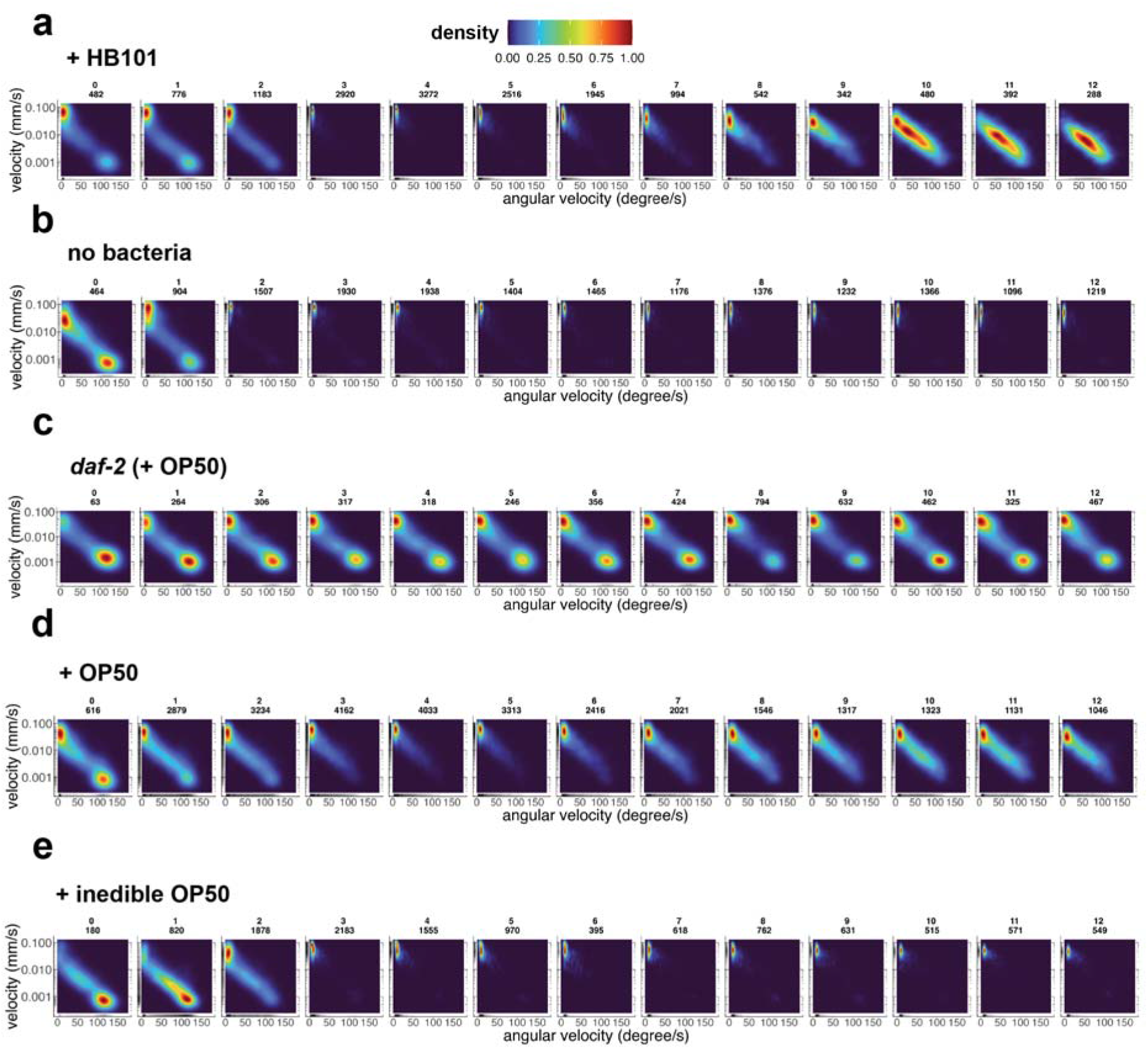
**(a-e)** Two-dimensional probability distributions of speed (mm/s) and angular velocity (degrees/s) averaged over 10 s time windows across the first 12 hours of dauer exit in different environments. Indicated is the time after transfer and the number of tracked paths within that time point. OP50, no bacteria and inedible OP50 are data from n=3 experiments, all other n=2.

**Supplemental figure S2.**
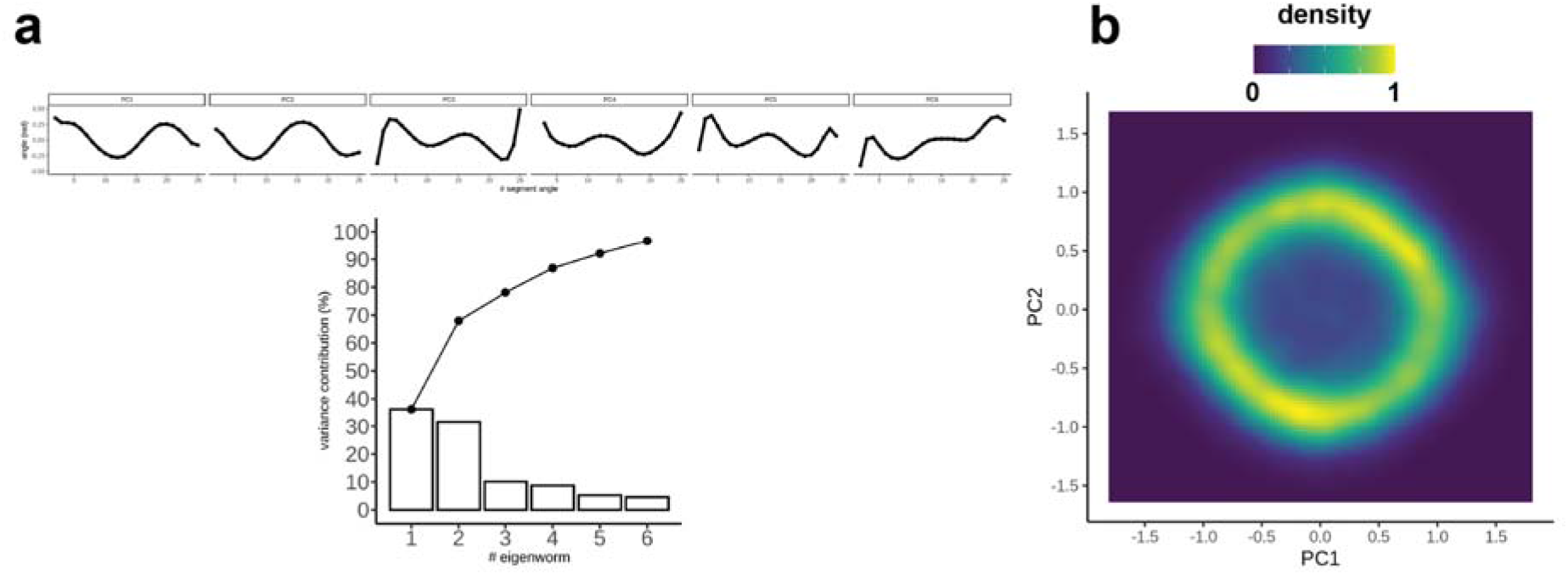
**(a)** Top 5 *Eigenworms* for moving worms in the +OP50 dataset across all time points (n=3). Bar plot shows the variance explained by each *Eigenworm.* Traces show cumulative values. **(b)** Probability density for the first two principal components.

**Supplemental figure S3.**
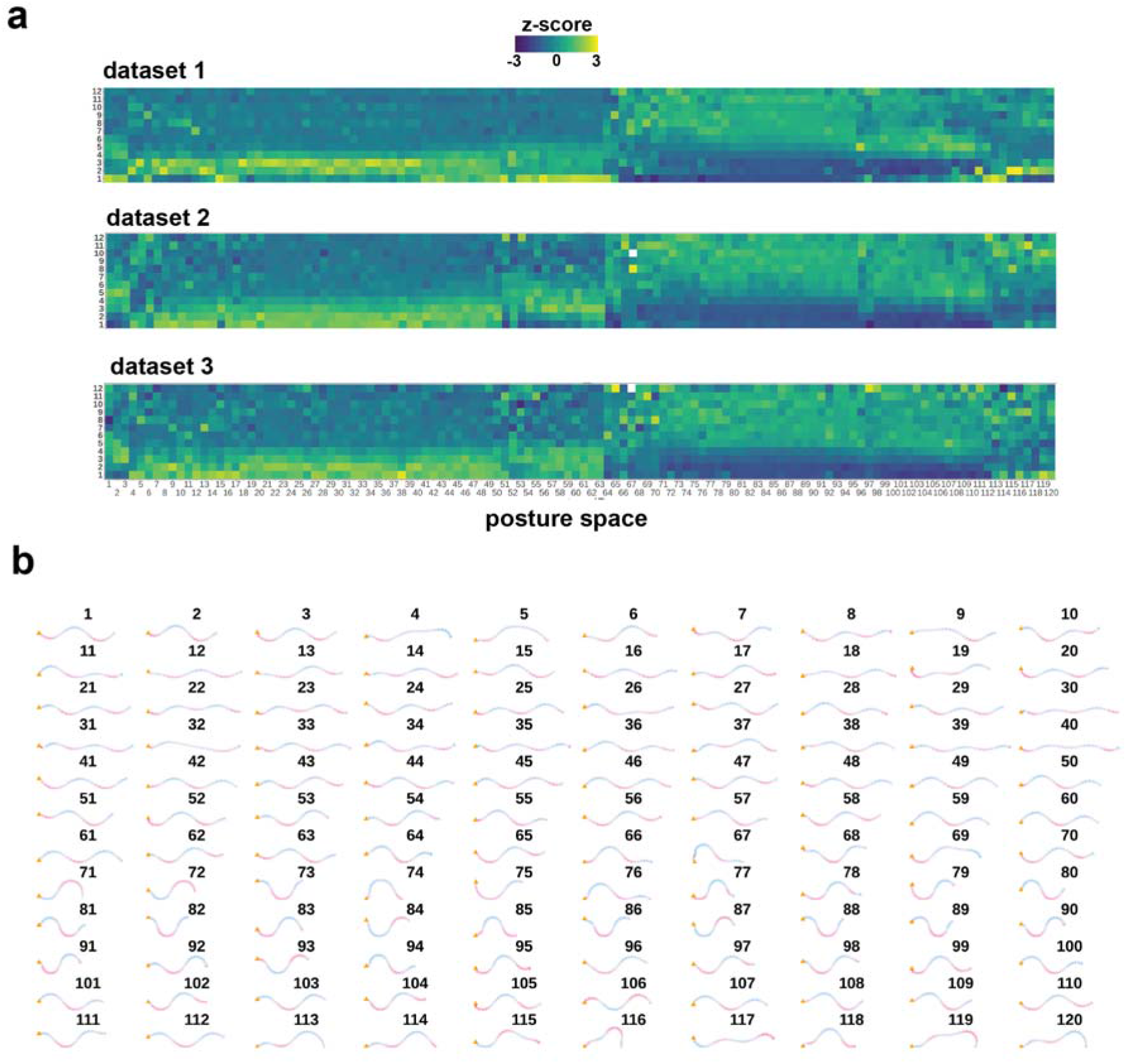
**(a)** Heatmaps of the relative occurrences of 120 postures during dauer exit on a food source (+ OP50), plotted individually but within the same posture space for 3 experimental repeats. Color indicates whether a posture is over or underrepresented at that time point relative to all time points. The postures that are corresponding to columns in the heatmaps are plotted in **(b)**.

**Supplemental figure S4.**
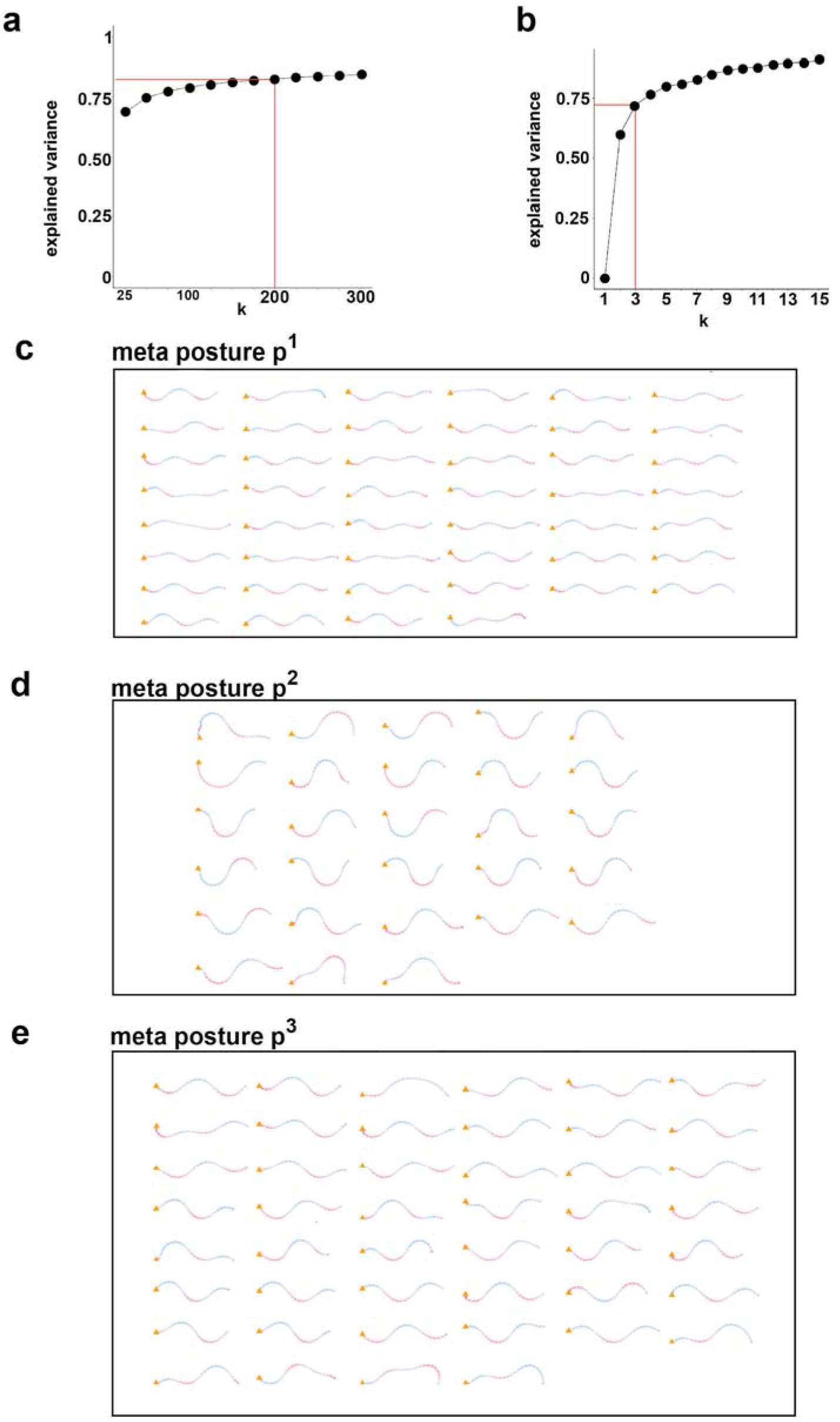
**(a)** Elbow plot for posture identification through k-means clustering of angle vectors of all skeletons, showing the explained variance with increasing k. **(b)** Same as (a), but for meta posture clustering, using the previously determined postures as input. Red line indicates the used k in our analysis. **(c-e)** Postures contained within the three meta posture groups.

**Supplemental figure S5.**
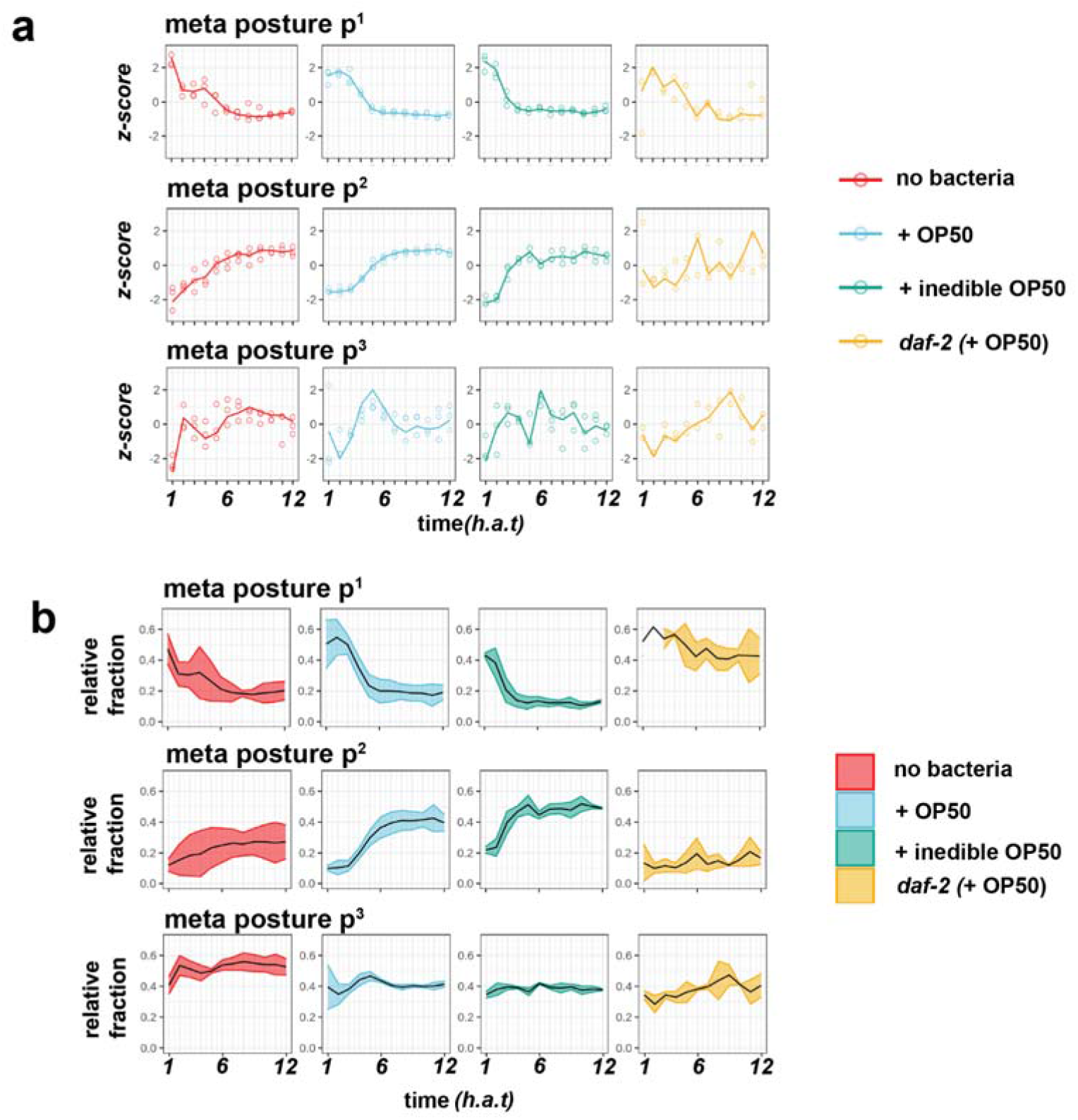
**(a)** Relative occurrence of the three meta postures during the first 12 hours of dauer exit, normalized to all time points within one experimental repeat. Each dot represents one experiment of n=3 experiments (except *daf-2,* n=2). **(b)** Same data as in (a), but plotting the relative abundance of each metaposture across time and all experimental repeats of one condition.

**Supplemental figure S6.**
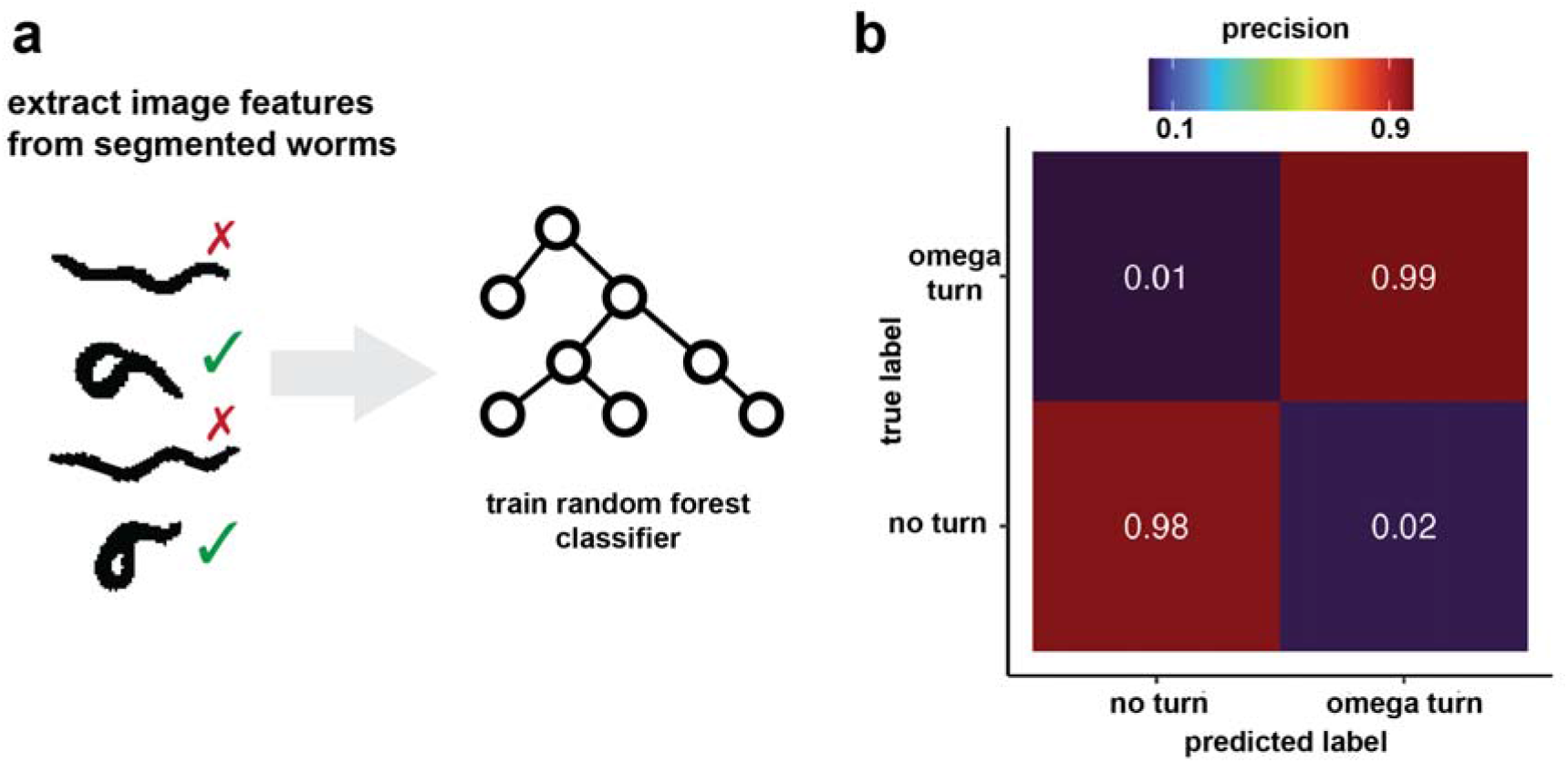
**(a)** Schematic of the training process for the random forest classifier. 17 image features are extracted from labeled worm bitmasks and used for training the random forest classifier in KNIME. For new data, prediction will occur for each segmented worm and frame. **(b)** Confusion matrix showing the classifier’s performance on a unseen test data set (93x omega turn, 89x “no turn”). Indicated is the precision score.

**Supplemental figure S7.**
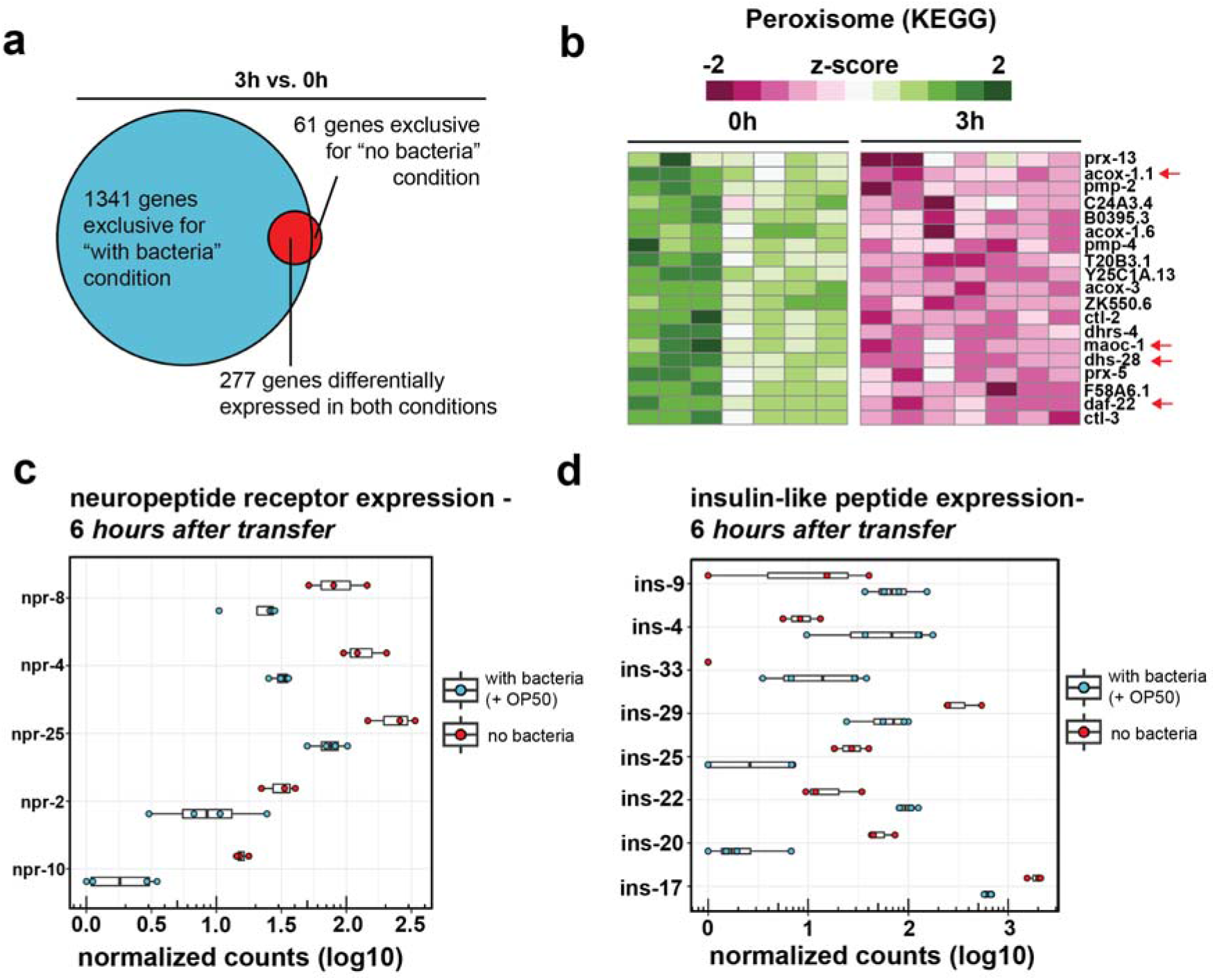
**(a)** Venn-diagram showing the overlap between differentially regulated genes when comparing 0h and 3h time points either in a condition with (blue) and without (red) a bacterial food source. **(b)** Genes within the KEGG peroxisome GO-term, the third most significant GO-term when using the list of 277 genes that are differentially expressed in both conditions as input. Highlighted are the members of the peroxisome beta-oxidation pathway. **(c)** Normalized counts of differentially expressed neuropeptide receptors when comparing both conditions at 6 *h.a.t.* **(d)** Normalized counts of differentially expressed insulin peptides when comparing both conditions at 6 *h.a.t.* All shown genes were significantly different when comparing the two conditions (Wald test, p-value adjusted <0.05).

## Bibliography

1. Kyle Honegger and Benjamin de Bivort. Stochasticity, individuality and behavior. Current Biology, 28(1):R8–R12, January 2018. doi: 10.1016/j.cub.2017.11.058.

2. J.G. White, E. Southgate, J.N. Thomson, and S. Brenner. The Structure of the Nervous System of the Nematode Caenorhabditis elegans. Philosophical Transactions of the Royal Society of London, 314(1165):1–340, 1986.

3. Steven W. Flavell and Andrew Gordus. Dynamic functional connectivity in the static connectome of Caenorhabditis elegans. Current Opinion in Neurobiology, 73:102515, April 2022. doi: 10.1016/j.conb.2021.12.002.

4. Randall C Cassada and Richard L Russell. The Dauerlarva, a Post-Embryonic Nematode Developmental elegans Variant of the Caenorhabditis. Developmental Biology, 342(46): 326–342, 1975.

5. Michael Klass and David Hirsh. Non-ageing developmental variant of Caenorhabditis elegans. Nature, 260(5551):523–525, 1976. doi: 10.1038/260523a0. ISBN: 0028-0836 (Print)$\backslash$r0028-0836 (Linking).

6. Nicole Fielenbach and Adam Antebi. C. elegans dauer formation and the molecular basis of plasticity. Genes and Development, 22(16):2149–2165, 2008. doi: 10.1101/gad.1701508. ISBN: 0890-9369 (Print).

7. L Ryan Baugh and Patrick J Hu. Starvation Responses Throughout the *Caenorhabditis elegans* Life Cycle. Genetics, 216(4):837–878, December 2020. doi: 10.1534/genetics.120.303565.

8. Patrice S Albert and Donald L Riddle. Developmental Alterations in Sensory Neuroanatomy of the Caenorhabditis elegans Dauer Larva. The Journal Of Comparative Neurology, 481: 461–481, 1983.

9. Nathan E Schroeder, Rebecca J Androwski, Alina Rashid, Harksun Lee, Junho Lee, and Maureen M Barr. Dauer-Specific Dendrite Arborization in C. elegans Is Regulated by KPC-1 / Furin. Current Biology, 23(16):1527–1535, 2013. doi: 10.1016/j.cub.2013.06.058. Publisher: Elsevier Ltd.

10. Abhishek Bhattacharya, Ulkar Aghayeva, Emily G. Berghoff, and Oliver Hobert. Plasticity of the Electrical Connectome of C. elegans. Cell, 0(0):1–16, January 2019. doi: 10.1016/j.cell.2018.12.024.

11. Sebastian Britz, Sebastian Matthias Markert, Daniel Witvliet, Anna Maria Steyer, Sarah Tröger, Ben Mulcahy, Philip Kollmannsberger, Yannick Schwab, Mei Zhen, and Christian Stigloher. Structural Analysis of the Caenorhabditis elegans Dauer Larval Anterior Sensilla by Focused Ion Beam-Scanning Electron Microscopy. Frontiers in Neuroanatomy, 15: 732520, November 2021. doi: 10.3389/fnana.2021.732520.

12. In Hae Lee, Carl Procko, Yun Lu, and Shai Shaham. Stress-Induced Neural Plasticity Mediated by Glial GPCR REMO-1 Promotes C. elegans Adaptive Behavior. Cell Reports, 34(2): 108607, 2021. doi: 10.1016/j.celrep.2020.108607. Publisher: ElsevierCompany.

13. Nathan Cermak, Stephanie K. Yu, Rebekah Clark, Yung Chi Huang, Saba N. Baskoylu, and Steven W. Flavell. Whole-organism behavioral profiling reveals a role for dopamine in state dependent motor program coupling in C. Elegans. eLife, 9:1–34, 2020. doi: 10.7554/eLife.57093.

14. Daniel Ramot, Brandon E. Johnson, Tommie L. Berry, Lucinda Carnell, and Miriam B. Goodman. The parallel worm tracker: A platform for measuring average speed and drug-induced paralysis in nematodes. PLoS ONE, 3(5):6–12, 2008. doi: 10.1371/journal.pone.0002208. ISBN: 1932-6203 (Electronic)$\backslash$n1932-6203 (Linking).

15. Eviatar Yemini, Tadas Jucikas, Laura J. Grundy, André E.X. Brown, and William R. Schafer. A database of Caenorhabditis elegans behavioral phenotypes. Nature Methods, 10(9):877–879, 2013. doi: 10.1038/nmeth.2560. ISBN: 1548-7105 (Electronic)$\backslash$r1548-7091 (Linking).

16. Shay Stern, Christoph Kirst, and Cornelia I. Bargmann. Neuromodulatory Control of Long-Term Behavioral Patterns and Individuality across Development. Cell, 0(0):1–14, 2017. doi: 10.1016/j.cell.2017.10.041. Publisher: Elsevier Inc.

17. Ida L. Barlow, Luigi Feriani, Eleni Minga, Adam McDermott-Rouse, Thomas James O’Brien, Ziwei Liu, Maximilian Hofbauer, John R. Stowers, Erik C. Andersen, Siyu Serena Ding, and André E. X. Brown. Megapixel camera arrays enable high-resolution animal tracking in multiwell plates. Communications Biology, 5(1):253, December 2022. doi: 10.1038/s42003-022-03206-1.

18. Jolle W. Jolles. Broad-scale applications of the Raspberry Pi: A review and guide for biologists. Methods in Ecology and Evolution, 12(9):1562–1579, September 2021. doi: 10.1111/2041-210X.13652.

19. Christian Dietz, Curtis T. Rueden, Stefan Helfrich, Ellen T. A. Dobson, Martin Horn, Jan Eglinger, Edward L. Evans, Dalton T. McLean, Tatiana Novitskaya, William A. Ricke, Nathan M. Sherer, Andries Zijlstra, Michael R. Berthold, and Kevin W. Eliceiri. Integration of the ImageJ Ecosystem in KNIME Analytics Platform. Frontiers in Computer Science, 2:8, March 2020. doi: 10.3389/fcomp.2020.00008.

20. Tobias Pietzsch, Stephan Preibisch, Pavel Tomančák, and Stephan Saalfeld. ImgLib2—generic image processing in Java. Bioinformatics, 28(22):3009–3011, November 2012. doi: 10.1093/bioinformatics/bts543.

21. Jean-Yves Tinevez, Nick Perry, Johannes Schindelin, Genevieve M. Hoopes, Gregory D. Reynolds, Emmanuel Laplantine, Sebastian Y. Bednarek, Spencer L. Shorte, and Kevin W. Eliceiri. TrackMate: An open and extensible platform for single-particle tracking. Methods, 115:80–90, February 2017. doi: 10.1016/j.ymeth.2016.09.016.

22. Marta Maria Gaglia, Cynthia Kenyon, and San Francisco. Stimulation of Movement in a Quiescent, Hibernation-Like Form of Caenorhabditis elegans by Dopamine Signaling. 29 (22):7302–7314, 2009. doi: 10.1523/JNEUROSCI.3429-08.2009.

23. Sreeparna Pradhan, Sabrina Quilez, Kai Homer, and Michael Hendricks. Environmental Programming of Adult Foraging Behavior in C. elegans. Current Biology, 29(17):2867–2879.e4, 2019. doi: 10.1016/j.cub.2019.07.045. Publisher: Elsevier Ltd.

24. Harksun Lee, Myung-kyu Choi, Daehan Lee, Hye-sung Kim, Hyejin Hwang, Heekyeong Kim, Sungsu Park, Young-ki Paik, and Junho Lee. Nictation, a dispersal behavior of the nematode Caenorhabditis elegans, is regulated by IL2 neurons. Nature Neuroscience, 15 (1), 2012. doi: 10.1038/nn.2975.

25. Antoine Barriere and Marie-Anne Felix. Natural variation and population genetics of Caenorhabditis elegans. WormBook: the online review of C. elegans biology, pages 1–19, 2005. doi: 10.1895/wormbook.1.43.1. ISBN: 1551-8507.

26. Marie-Anne Félix and Fabien Duveau. Population dynamics and habitat sharing of natural populations of Caenorhabditis elegans and C. briggsae. BMC Biology, 10(1):59, December 2012. doi: 10.1186/1741-7007-10-59.

27. Wolkow, C.A. and Hall, D.H. The Dauer Muscle. In WormAtlas. 2013.

28. Juliette Ben Arous, Sophie Laffont, and Didier Chatenay. Molecular and sensory basis of a food related two-state behavior in C. elegans. PLoS ONE, 4(10):1–8, 2009. doi: 10.1371/journal.pone.0007584.

29. Greg J. Stephens, Bethany Johnson-Kerner, William Bialek, and William S. Ryu. Dimensionality and dynamics in the behavior of C. elegans. PLoS Computational Biology, 4(4), 2008. doi: 10.1371/journal.pcbi.1000028.

30. André E.X. Brown, Eviatar I. Yemini, Laura J. Grundy, Tadas Jucikas, and William R. Schafer. A dictionary of behavioral motifs reveals clusters of genes affecting Caenorhabditis elegans locomotion. Proceedings of the National Academy of Sciences of the United States of America, 110(2):791–796, 2013. doi: 10.1073/pnas.1211447110.

31. Bertalan Gyenes and André E.X. Brown. Deriving shape-based features for C. elegans locomotion using dimensionality reduction methods. Frontiers in Behavioral Neuroscience, 10(AUG):1–9, 2016. doi: 10.3389/fnbeh.2016.00159.

32. Ingrid Hums, Julia Riedl, Fanny Mende, Saul Kato, Harris S Kaplan, Richard Latham, Michael Sonntag, Lisa Traunmüller, and Zimmer, Manuel. Regulation of two motor patterns enables the gradual adjustment of locomotion strategy in Caenorhabditis elegans. eLife, 5 (e14116):1–36, 2016. doi: 10.7554/eLife.14116.

33. Harris S Kaplan, Oriana Salazar Thula, Niklas Khoss, and Manuel Zimmer. Nested neuronal dynamics orchestrate a behavioral hierarchy across timescales. Neuron, pages 1–15, 2019. doi: 10.1016/j.neuron.2019.10.037. Publisher: Elsevier Inc.

34. Roland F. Schwarz, Robyn Branicky, Laura J. Grundy, William R. Schafer, and André E.X. Brown. Changes in Postural Syntax Characterize Sensory Modulation and Natural Variation of C. elegans Locomotion. PLoS Computational Biology, 11(8):1–16, 2015. doi: 10.1371/journal.pcbi.1004322. ISBN: 1553-7404 (Electronic)$\backslash$n1553-7390 (Linking).

35. Jesse M Gray, Joseph J Hill, and Cornelia I Bargmann. A circuit for navigation in Caenorhabditis elegans. 2005.

36. Sreekanth H. Chalasani, Nikos Chronis, Makoto Tsunozaki, Jesse M. Gray, Daniel Ramot, Miriam B. Goodman, and Cornelia I. Bargmann. Dissecting a circuit for olfactory behaviour in Caenorhabditis elegans. Nature, 450(7166):63–70, 2007. doi: 10.1038/nature06292. ISBN: 1476-4687 (Electronic)$\backslash$r0028-0836 (Linking).

37. Patrick Laurent, Zoltan Soltesz, Geoff Nelson, Changchun Chen, Fausto Arellano-Carbajal, Emmanuel Levy, and Mario de Bono. Decoding a neural circuit controlling global animal state in C. Elegans. eLife, 2015(4):1–39, 2015. doi: 10.7554/eLife.04241.

38. Steven W Flavell, Navin Pokala, Evan Z Macosko, Dirk R Albrecht, Johannes Larsch, and Cornelia I Bargmann. Serotonin and the Neuropeptide PDF Initiate and Extend Opposing Behavioral States in C. elegans. Cell, 154(5):1023–1035, 2013. doi: 10.1016/j.cell.2013.08.001. Publisher: Elsevier Inc.

39. Berta Vidal, Ulkar Aghayeva, Haosheng Sun, Chen Wang, Lori Glenwinkel, Emily A. Bayer, and Oliver Hobert. An atlas of Caenorhabditis elegans chemoreceptor expression. PLOS Biology, 16(1):e2004218, January 2018. doi: 10.1371/journal.pbio.2004218. ISBN: 1111111111.

40. Sarah B. Pierce, Michael Costa, Robert Wisotzkey, Sharmila Devadhar, Sheila A. Homburger, Andrew R. Buchman, Kimberly C. Ferguson, Jonathan Heller, Darren M. Platt, Amy A. Pasquinelli, Leo X. Liu, Stephen K. Doberstein, and Gary Ruvkun. Regulation of DAF-2 receptor signaling by human insulin and *ins-1*, a member of the unusually large and diverse *C. elegans* insulin gene family. Genes & Development, 15(6):672–686, March 2001. doi: 10.1101/gad.867301.

41. A. Cornils, M. Gloeck, Z. Chen, Y. Zhang, and J. Alcedo. Specific insulin-like peptides encode sensory information to regulate distinct developmental processes. Development, 138(6):1183–1193, 2011. doi: 10.1242/dev.060905. ISBN: 1477-9129 (Electronic)$\backslash$r0950-1991 (Linking).

42. Weiqing Li, Scott G. Kennedy, and Gary Ruvkun. daf-28 encodes a C. elegans insulin superfamily member that is regulated by environmental cues and acts in the DAF-2 signaling pathway. Genes and Development, 17(7):844–858, 2003. doi: 10.1101/gad.1066503. ISBN: 0890-9369 (Print)$\backslash$n0890-9369 (Linking).

43. Tamar Hashimshony, Florian Wagner, Noa Sher, and Itai Yanai. CEL-Seq: Single-Cell RNA-Seq by Multiplexed Linear Amplification. Cell Reports, 2(3):666–673, 2012. doi: 10.1016/j.celrep.2012.08.003. ISBN: 2211-1247 (Electronic) Publisher: The Authors.

44. Stephan H. Von Reuss, Neelanjan Bose, Jagan Srinivasan, Joshua J. Yim, Joshua C. Judkins, Paul W. Sternberg, and Frank C. Schroeder. Comparative metabolomics reveals biogenesis of ascarosides, a modular library of small-molecule signals in C. elegans. Journal of the American Chemical Society, 134(3):1817–1824, 2012. doi: 10.1021/ja210202y.

45. W. L. Hung, Y. Wang, J. Chitturi, and M. Zhen. A Caenorhabditis elegans developmental decision requires insulin signaling-mediated neuron-intestine communication. Development, 141(8):1767–1779, 2014. doi: 10.1242/dev.103846.

46. James W. Golden and Donald L. Riddle. A Caenorhabditis elegans dauer-inducing pheromone and an antagonistic component of the food supply. Journal of Chemical Ecology, 10(8):1265–1280, 1984. doi: 10.1007/BF00988553. ISBN: 0098-0331.

47. Umer Saleem Bhat, Navneet Shahi, Siju Surendran, and Kavita Babu. Neuropeptides and Behaviors: How Small Peptides Regulate Nervous System Function and Behavioral Outputs. Frontiers in Molecular Neuroscience, 14:786471, December 2021. doi: 10.3389/fnmol.2021.786471.

48. Yohei Matsunaga, Kensuke Nakajima, Keiko Gengyo-Ando, Shohei Mitani, Takashi Iwasaki, and Tsuyoshi Kawano. A caenorhabditis elegans insulin-like peptide, INS-17: Its physiological function and expression pattern. Bioscience, Biotechnology and Biochemistry, 76(11): 2168–2172, 2012. doi: 10.1271/bbb.120540.

49. Mochi Liu, Sandeep Kumar, Anuj K. Sharma, and Andrew M. Leifer. A high-throughput method to deliver targeted optogenetic stimulation to moving C. elegans populations. PLOS Biology, 20(1):e3001524, January 2022. doi: 10.1371/journal.pbio.3001524.

50. Laetitia Hebert, Tosif Ahamed, Antonio C. Costa, Liam O’Shaughnessy, and Greg J. Stephens. WormPose: Image synthesis and convolutional networks for pose estimation in C. elegans. PLoS Computational Biology, 17(4):1–20, 2021. doi: 10.1371/journal.pcbi.1008914. ISBN: 1111111111.

51. Avelino Javer, Michael Currie, Chee Wai Lee, Jim Hokanson, Kezhi Li, Céline N. Martineau, Eviatar Yemini, Laura J. Grundy, Chris Li, QueeLim Ch’ng, William R. Schafer, Ellen A. A. Nollen, Rex Kerr, and André E. X. Brown. An open-source platform for analyzing and sharing worm-behavior data. Nature Methods, 15(9):645–646, 2018. doi: 10.1038/s41592-018-0112-1. Publisher: Springer US.

52. Lise Frézal and Marie-Anne Félix. C. elegans outside the Petri dish. eLife, 4:e05849, March 2015. doi: 10.7554/eLife.05849.

53. Scott J. Dixon, Mariam Alexander, Kevin Ka Ming Chan, and Peter John Roy. Insulinlike signaling negatively regulates muscle arm extension through DAF-12 in Caenorhabditis elegans. Developmental Biology, 318(1):153–161, 2008. doi: 10.1016/j.ydbio.2008.03.019. ISBN: 1095-564X (Electronic).

54. Cornelia I Bargmann and H Robert Horvitz. Control of Larval Development by Chemosensory Neurons in Caenorhabditis elegans. Science, 17(1972), 1991.

55. Gáspár Jékely. Global view of the evolution and diversity of metazoan neuropeptide signaling. Proceedings of the National Academy of Sciences, 110(21):8702–8707, May 2013. doi: 10.1073/pnas.1221833110.

56. Arantza Barrios, Rajarshi Ghosh, Chunhui Fang, Scott W Emmons, and Maureen M Barr. PDF-1 neuropeptide signaling modulates a neural circuit for mate-searching behavior in C. elegans. Nature Neuroscience, 15(12):1675–1682, December 2012. doi: 10.1038/nn.3253.

57. Michal Turek, Judith Besseling, Jan-Philipp Spies, Sabine König, and Henrik Bringmann. Sleep-active neuron specification and sleep induction require FLP-11 neuropeptides to systemically induce sleep. eLife, 5:e12499, March 2016. doi: 10.7554/eLife.12499.

58. James Siho Lee, Pei-Yin Shih, Oren N. Schaedel, Porfirio Quintero-Cadena, Alicia K. Rogers, and Paul W. Sternberg. FMRFamide-like peptides expand the behavioral repertoire of a densely connected nervous system. Proceedings of the National Academy of Sciences, 114(50):E10726–E10735, December 2017. doi: 10.1073/pnas.1710374114.

59. Cynthia M. Chai, Mahdi Torkashvand, Maedeh Seyedolmohadesin, Heenam Park, Vivek Venkatachalam, and Paul W. Sternberg. Interneuron control of C. elegans developmental decision-making. Current Biology, page S0960982222005644, April 2022. doi: 10.1016/j.cub.2022.03.077.

60. Mi Cheong Cheong, Alexander B. Artyukhin, Young Jai You, and Leon Avery. An opioid-like system regulating feeding behavior in C. elegans. eLife, 4:1–19, 2015. doi: 10.7554/eLife.06683.

61. Mykola Mylenko, Sebastian Boland, Sider Penkov, Julio L. Sampaio, Benoit Lombardot, Daniela Vorkel, Jean Marc Verbavatz, and Teymuras V. Kurzchalia. NAD+ is a food component that promotes exit from dauer diapause in Caenorhabditis elegans. PLoS ONE, 11 (12):1–17, 2016. doi: 10.1371/journal.pone.0167208.

62. Junyue Cao, Jonathan S Packer, Vijay Ramani, Darren A Cusanovich, Chau Huynh, Riza Daza, Xiaojie Qiu, Choli Lee, Scott N Furlan, Frank J Steemers, Andrew Adey, Robert H Waterston, Cole Trapnell, and Jay Shendure. Comprehensive single-cell transcriptional profiling of a multicellular organism. Science, 357(6352):661–667, August 2017. doi: 10.1126/science.aam8940.

63. Seth R. Taylor, Gabriel Santpere, Alexis Weinreb, Alec Barrett, Molly B. Reilly, Chuan Xu, Erdem Varol, Panos Oikonomou, Lori Glenwinkel, Rebecca McWhirter, Abigail Poff, Manasa Basavaraju, Ibnul Rafi, Eviatar Yemini, Steven J. Cook, Alexander Abrams, Berta Vidal, Cyril Cros, Saeed Tavazoie, Nenad Sestan, Marc Hammarlund, Oliver Hobert, and David M. Miller. Molecular topography of an entire nervous system. Cell, 184(16):4329–4347.e23, August 2021. doi: 10.1016/j.cell.2021.06.023.

64. Karoline Holler and Jan Philipp Junker. RNA tomography for spatially resolved transcriptomics (tomo-seq). Methods in Molecular Biology, 1920:129–141, 2019. doi: 10.1007/978-1-4939-9009-2_9. ISBN:9781493990092.

65. Alexander Dobin, Carrie A. Davis, Felix Schlesinger, Jorg Drenkow, Chris Zaleski, Sonali Jha, Philippe Batut, Mark Chaisson, and Thomas R. Gingeras. STAR: Ultrafast universal RNA-seq aligner. Bioinformatics, 29(1):15–21, 2013. doi: 10.1093/bioinformatics/bts635. ISBN: 1367-4811 (Electronic)\n1367-4803 (Linking).

66. Yang Liao, Gordon K Smyth, and Wei Shi. The R package Rsubread is easier, faster, cheaper and better for alignment and quantification of RNA sequencing reads. Nucleic Acids Research, 47(8):e47–e47, May 2019. doi: 10.1093/nar/gkz114.

67. Michael I Love, Wolfgang Huber, and Simon Anders. Moderated estimation of fold change and dispersion for RNA-seq data with DESeq2. Genome Biology, 15(12):550, December 2014. doi: 10.1186/s13059-014-0550-8.

68. Liis Kolberg, Raudvere, Uku, Kuzmin, Ivan, Vilo, Jaak, and Peterson, Hedi. gprofiler2 – an R package for gene list functional enrichment analysis and namespace conversion toolset g:Profiler. F1000Research, 9(ELIXIR):709, 2020. doi: https://doi.org/10.12688/f1000research.24956.2.

